# Microbial community assembly in a multi-layer dendritic metacommunity

**DOI:** 10.1101/2020.01.31.929562

**Authors:** Nathan I. Wisnoski, Jay T. Lennon

## Abstract

A major goal of metacommunity ecology is to infer the local- and regional-scale processes that underlie community assembly. In dendritic ecological networks (e.g., stream metacommunities), branching and directional dispersal connectivity can alter the balance between local and regional factors during assembly. However, the implications of vertical habitat structure (e.g., planktonic versus benthic sediments) in dendritic metacommunities remain unclear. In this study, we analyzed the bacterial metacommunity of a fifth-order mountain stream network to assess habitat differences in the (1) dominant community assembly processes, (2) spatial scaling of community assembly processes, and (3) longitudinal variation in community assembly. Using taxonomic and phylogenetic null modeling approaches, we found habitat-specific spatial patterns of community assembly across the dendritic network. Compositional differences between planktonic and benthic communities were maintained by divergent species sorting, but stochasticity influenced assembly at local scales. Planktonic communities showed scale-dependent assembly, transitioning from convergent sorting at local scales to divergent sorting at regional scales, while sediment community assembly was less scale dependent (convergent sorting remained important across all scales). While divergent sorting structured headwaters in both habitat types, sediment communities converged in structure downstream. Taken together, our results show that vertical habitat structure regulates the scale-dependent processes of community assembly across the dendritic metacommunity.

## INTRODUCTION

Metacommunity ecology examines the assembly, structure, and diversity of communities with an emphasis on the interplay between local- and regional-scale processes (Leibold and Chase 2018). At the local scale, environmental filtering and species interactions influence assembly through species sorting (Leibold 1998, Chase et al. 2005), which leads to similar communities in similar habitats (i.e., convergent species sorting) and dissimilar communities in dissimilar habitats (i.e., divergent species sorting). The metacommunity framework also incorporates the effects of dispersal and stochastic processes on community assembly (Mouquet and Loreau 2003, Zhou and Ning 2017). For example, dispersal limitation can account for compositional dissimilarity between communities in similar habitats, while rampant dispersal can homogenize communities across dissimilar habitats due to mass effects. Therefore, species sorting should play a prevailing role in structuring communities when dispersal is low but non-limiting.

While the direction of dispersal is often assumed to be random in metacommunities, some ecosystems have physical features that impose directionality. For example, stream and river ecosystems represent dendritic networks with hierarchical, branching connectivity that constrains and directionally orients dispersal (Fig. 1A) (Grant et al. 2007, Brown et al. 2011, Carrara et al. 2012, Altermatt 2013). As a result, some sites in dendritic networks are more isolated and less connected than others. For example, headwater streams are separated by elongated dispersal routes along the stream network that may exceed dispersal capabilities of some organisms. At the same time, dispersal is counteracted by prevailing downstream flows that further reduce headwater connectivity with the metacommunity (Brown et al. 2011, Altermatt 2013, Tonkin et al. 2018). Many headwater communities (e.g., benthic macroinvertebrates) are assembled by species sorting, while downstream communities show greater environmental mismatch due to high rates of dispersal from upstream (i.e., mass effects) (Brown and Swan 2010, Tornwall et al. 2017). However, different patterns have been documented for other taxonomic groups with limited upstream-dispersal vectors, such as passively dispersing microorganisms. For these communities, headwater assemblages experience high rates of immigration from surrounding terrestrial ecosystems that can disrupt species sorting (Ruiz-González et al. 2015, Battin et al. 2016). Terrestrial-derived bacteria are gradually filtered out as they disperse downstream, where species sorting becomes the dominant process as stable planktonic communities establish in reaches with longer residence times (Read et al. 2015, Savio et al. 2015, Ruiz-González et al. 2015).

**Fig. 1.**
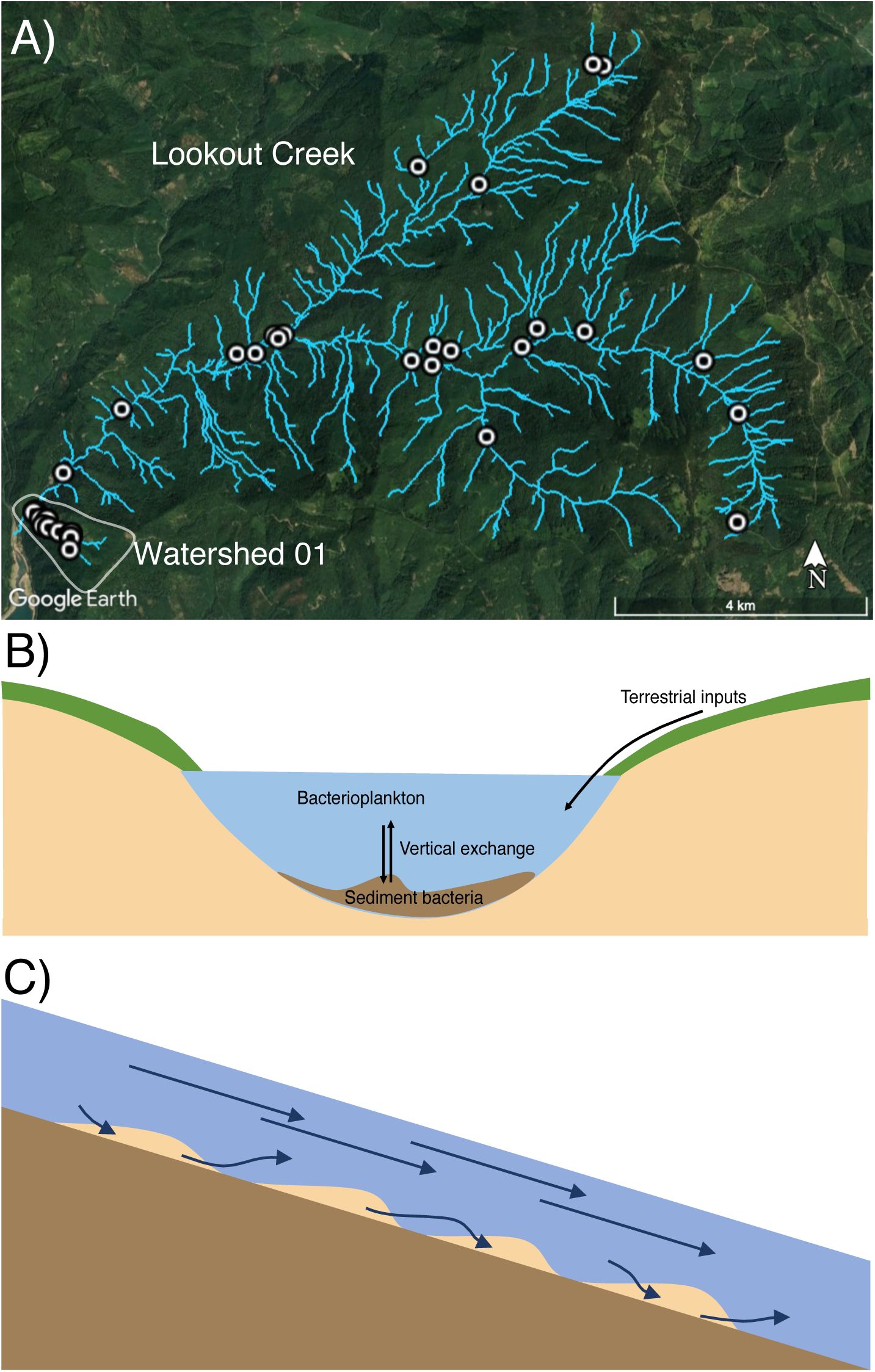
The dendritic metacommunity structure of stream ecosystems with vertical habitat structure. (A) Map of sampling locations within H. J. Andrews Experimental Forest. Sampling was conducted extensively across the broader Lookout Creek watershed, spanning stream orders 1 to 5. Sampling was also conducted intensively within small Watershed 1 (lower left). Imagery sourced from Google Earth Pro, with stream network sourced from H.J. Andrews Experimental Forest data portal. (B) A lateral cross-section through the stream channel reveals the vertical habitat structure that is present in streams. Bacterioplankton occur in the water column, while sediment-attached biofilms line the benthic habitat. (C) A longitudinal cross-section of the stream channel demonstrates the differences in spatial connectivity between planktonic and benthic habitats, where plankton are hypothesized to have higher dispersal than sediment-attached bacteria.

Another feature of dendritic systems that is not considered by classical metacommunity theory is that they commonly exhibit vertical habitat structure (Fig. 1B). In streams, planktonic organisms inhabiting the water column experience vastly different physical environments than benthic organisms living in the sediment matrix of the streambed (Hart and Finelli 1999). As a result, different sets of environmental filters may influence the composition of planktonic and benthic bacterial communities (Besemer et al. 2012, Wilhelm et al. 2013). For example, planktonic microorganisms must contend with changes in resource availability, pH, predation, and hydrology (Fierer et al. 2007, Read et al. 2015, Niño-García et al. 2016), while benthic communities experience additional constraints, such as shear stress, space limitation in biofilms, and fluctuating redox conditions resulting from surface water-groundwater mixing (Battin et al. 2016). The different flow environments of benthic and planktonic habitats could also affect bacterial dispersal rates and community assembly (Battin et al. 2016). For example, bacterioplankton presumably have high dispersal rates that increase the potential for mass effects or stochastic processes, while bacteria in sediment biofilms disperse downstream intermittently (Leff et al. 1992), increasing the potential for species sorting (Fig. 1C). However, the two habitats are not completely separate, as planktonic-benthic mixing introduces a vertical axis of dispersal allowing plankton to colonize sediments and sediment-associated bacteria to be suspended in the water column (Leff et al. 1992, Freimann et al. 2015), which may influence community structure at local scales. These habitat-specific differences in environmental filters and dispersal could alter the relative importance of community assembly processes underlying local and regional diversity by influencing their spatial distributions in the dendritic network.

In this study, we analyzed bacterial diversity in a dendritic metacommunity while considering not only directional flow, but also the vertical habitat structure that influences dispersal between stream sediments and the overlying water column, as well as the rate of downstream movement through the network. Specifically, we analyzed planktonic and sediment-associated bacterial communities in a mountain stream network. Using taxonomic and phylogenetic approaches, we tested whether the relative importance of community assembly processes varied (1) between planktonic and benthic habitats, (2) across spatial scales, and (3) along the longitudinal (i.e., headwater versus downstream) stream dimension in the dendritic metacommunity.

## METHODS

### Study Site

H.J. Andrews Experimental Forest (44.2° N, 122.2° W) is a 6,400-hectare conifer forest in the Western Cascade Range, Oregon, USA. Andrews Forest is a Long-Term Ecological Research (LTER) site that contains the Lookout Creek watershed, a fifth-order, mountainous (410–1630 m elevation) catchment of high gradient streams that drains to the McKenzie River (Fig. 1). The underlying geology is volcanic and dates back to the Oligocene, with Miocene-age andesite lava flows at higher elevations (Swanson and James 1975). Catchment topography is steep with confined valleys, and precipitation filters through loamy, organic soils to the stream (Harr 1977). Streams are boulder-dominated, with step–pool, riffle–pool, and cascade reaches. At lower elevations, vegetation is primarily Douglas fir (*Pseudotsuga menziesii*), western hemlock (*Tsuga heterophylla*), and western red cedar (*Thuja plicata*). Pacific silver fir (*Abies amabilis*) and noble fir (*Abies procera*) are present at higher elevations. The climate is Mediterranean, with peak precipitation between October and April. Mean annual precipitation is 230 cm at low elevations and 355 cm at high elevations (McKee and Bierlmaier 1987), and a sizeable snowpack accumulates above 900 m (Daly et al. 2010).

### Sampling

In June 2015, we sampled streams in the Lookout Creek watershed of H.J. Andrews Experimental Forest (Fig. 1). Our sampling design was hierarchical, such that lower-order stream sites were nested within branches of higher-order stream sites. Our samples spanned all five stream orders of Lookout Creek, where headwaters are 1^st^-order streams. We sampled major confluences across the catchment. Each sampling location was geo-referenced using handheld GPS. At each site, we measured temperature, pH, and conductivity in the stream using a YSI 6920 V2-2 water quality sonde (YSI Incorporated, Yellow Springs, OH). We preserved water samples with HCl to pH 2 for chemical analyses in the laboratory. With the preserved water samples, we measured total nitrogen (TN) after persulfate digestion using the second derivative method (Bachmann and Canfield 1996) and total phosphorus (TP) using the ammonium molybdate method (Prepas and Rigler 1982). Dissolved organic carbon (DOC) was measured in 0.7-µm glass fiber filtered samples by oxidation and nondispersive infrared detection on a Shimadzu TOC-V (Kyoto, Japan). These environmental variables were used to capture longitudinal patterns in environmental conditions in the stream network.

To characterize bacterial communities, we sampled planktonic and sediment-associated microbial biomass for high-throughput community sequencing at each site. We sampled planktonic microorganisms by filtering 1 L of surface water onto 47 mm 0.2-µm Supor Filters (Pall, Port Washington, NY) in the field. We sampled sediment-associated communities (of sediment grain < 1 cm in diameter) using a sediment corer. All samples were frozen on dry ice in the field and preserved at –20 °C until processing. In the laboratory, we detached bacterial cells from sediment biofilms by gently sonicating 5 g of sediment in a 1% tetrasodium pyrophosphate solution for 10 min in pulses of 10 sec on, 5 sec off. We then used the cell suspension for downstream analysis of the sediment-associated community.

### Sequence Preparation and Processing

We characterized bacterial community composition by sequencing the 16S rRNA gene (Caporaso et al. 2012). We extracted DNA from surface water samples using the PowerWater DNA isolation kit (MoBio, Carlsbad, CA) and from the sediment extractions using the PowerSoil DNA isolation kit (MoBio, Carlsbad, CA). We PCR-amplified the V4 region of the 16S rRNA gene using barcoded primers (515F and 806R) for the Illumina MiSeq platform. Per each 50 µl reaction, PCR conditions were the following: 5 µl of 10X Perfect Taq Plus PCR Buffer (5Prime), 10 µl 5P solution (5Prime), 0.25 µl Perfect Taq Plus DNA Polymerase (5Prime), 1 µl dNTP mix (10 mM each), 1 µl 515F forward primer (10 µM), 1 µl 806R reverse primer (10 µM), and 10 ng of template. Thermal cycler conditions were 3 min at 94 °C, 30 cycles of (45 sec at 94 °C, 30 sec at 50 °C, and 90 sec at 72 °C), then 10 min at 72 °C. Sequence libraries were cleaned using AMPure XP purification kit, quantified using Quant-iT PicoGreen dsDNA assay kit (Invitrogen), and pooled at equal concentrations of 10 ng per library. We sequenced the pooled libraries on the Illumina MiSeq platform at the Indiana University Center for Genomics and Bioinformatics using 300 × 300 bp paired end reads (600-cycle Reagent Kit v3). We processed the raw reads using *mothur* to remove non-bacterial sequences and low-quality reads, and removed chimeras with VSEARCH (Schloss et al. 2009, Rognes et al. 2016). We classified OTUs with the OptiClust algorithm (Westcott and Schloss 2017) based on 97% similarity using the SILVA rRNA database version 132 (Quast et al. 2013). All further analyses were conducted in R version 3.5.3 (R Core Team 2018).

### Diversity Analyses

We analyzed taxonomic patterns of diversity within and between planktonic and benthic sediment habitats in the metacommunity. First, we rarefied each sample to a total number of 10,623 reads (the smallest sample with > 10,000 reads), and relativized reads for each OTU to the size of each sample using the *vegan* R package (Oksanen et al. 2019). As a measure of within-site (α) diversity, we used the exponential of Shannon’s index, which corresponds to the number of equally abundant species needed to obtain the value of Shannon diversity obtained on the original data (Jost 2007). To measure differences in community structure among sites (β-diversity), we calculated pairwise dissimilarities between communities using the Hellinger distance (Legendre and Gallagher 2001). To determine whether β-diversity was related to qualitative features of the stream network, such as habitat, stream order, and watershed, we used PERMANOVA (Anderson 2001). We used redundancy analysis (RDA) to quantify the importance of quantitative environmental variables (TP, TN, DOC, pH, elevation, conductivity) for explaining β-diversity (Legendre and Legendre 2012). We used multiple regression to quantify how community dissimilarity changed with increasing dendritic distance (i.e., along the stream network path) between sites within and between habitat types. We calculated dendritic distances in Google Earth using GIS layers of the H.J. Andrews stream network created from LIDAR imaging.

### Community assembly processes

We used a null model approach to distinguish deterministic species sorting from stochastic assembly processes across the stream network (Chase et al. 2011, Chase and Myers 2011, Stegen et al. 2015). In this approach, we used taxonomic and phylogenetic information from the bacterial sequencing efforts. Phylogenies organize bacterial taxa by their evolutionary history and can inform mechanisms of community assembly if broad-scale, ecologically relevant traits map onto phylogenetic relatedness (Cadotte and Davies 2016). Thus, environments may select phylogenetically similar subsets of taxa from the metacommunity that possess traits necessary to colonize the local habitat through species sorting. Convergent species sorting (i.e., communities favoring similar taxa due to environmental similarities) was inferred when pairwise phylogenetic β-diversity was lower than expected under stochastic assembly. In contrast, divergent species sorting (i.e., dissimilar environments favoring dissimilar taxa) was inferred when phylogenetic β-diversity was greater than stochastic expectations.

To calculate phylogenetic β-diversity, we first created a phylogeny of all the OTUs in the stream network using a double-precision, approximately maximum-likelihood approach with the program FastTree v. 2.1.8 (Price et al. 2010). Using the *picante* R package (Kembel et al. 2010), we computed the β-Mean Nearest Taxon Distance (βMNTD), an abundance-weighted community-scale measure of the mean phylogenetic relatedness of each OTU within a community compared to its most closely related OTU in a second community. We generated null distributions (n = 999) of βMNTD by randomly shuffling the tips of the phylogenetic tree. Because the contribution of rare taxa to βMNTD is small yet computationally intensive, we performed this analysis using only the OTUs detected at least 10 times in the metacommunity (n ≈ 5,700). For each pair of sites *i* and *j*, we then compared the observed βMNTD values to the null distribution for the site-pair to calculate the β-Mean Nearest Taxon Index (βNTI), which quantifies the degree of phylogenetic turnover relative to expected turnover under stochastic community assembly:

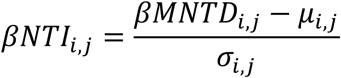

where *βMNTD*_*i,j*_ is the observed mean nearest taxon distance and the null distribution is described by its mean (*μ*_*i,j*_) and variance (*σ*_*i,j*_). Thus, βNTI is a z-score, and deviations are considered significant if |*βNTI*| > 2, where values greater than 2 indicate divergent sorting and values less than –2 indicate convergent sorting.

To test for stochastic assembly in sites with non-significant βNTI values (i.e., weak sorting), we compared observed taxonomic β-diversity to expectations generated by a stochastically assembled null model. For a pair of sites, high dispersal should decrease β-diversity from stochastic expectations, but dispersal limitation should increase β-diversity (Chase et al. 2011, Chase and Myers 2011). To quantify the contributions of these two processes, we modified the abundance-based Raup-Crick approach of Stegen et al. (2015) to generate distributions of expected dissimilarity values for each site-pair using the Hellinger distance (n = 999 permutations). The stochastic assembly null model was performed in the following way: OTUs were randomly selected in proportion to their regional site incidence, individuals were then sequentially and randomly added to local communities in proportion to their regional relative abundances, and total abundances of assembled communities were constrained to match observed total abundances. For each pair of sites, observed Hellinger distance was compared to the site-specific null distribution to compute *β*_*RC, Hellinger*_:

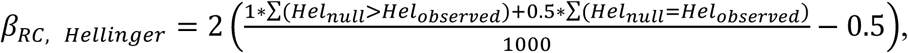

where ∑(*Hel*_*null*_ > *Hel*_*observed*_) is the number of null Hellinger distances greater than observed values and ∑(*Hel*_*null*_ = *Hel*_*observed*_) is the number of ties. After this calculation, *β*_*RC,Hellinger*_ ranges from −1 to 1. Deviations from null expectation were inferred when |*β* _*RC,Hellinger*_ | > 0.95, with *β*_*RC,Hellinger*_ > 0.95 indicating possible dispersal limitation and *β*_*RC,Hellinger*_ < −0.95 indicating potential mass effects.

### Scale-dependent and longitudinal patterns of assembly

Finally, we investigated whether the relative importance of community assembly processes varied across spatial scales and along the longitudinal axis of the stream network. When assessing the scale-dependence of community assembly processes in the dendritic metacommunity, we only compared sites that were hydrologically connected by flow (i.e., hierarchical upstream-downstream linkages but not among hydrologically disconnected headwaters). We calculated the dendritic distance separating each pair of sites, rounding distances to the nearest log_10_(m) to generate discrete distance classes spanning five orders of magnitude. We calculated the proportion of each assembly mechanism inferred within each distance class and quantified the frequencies of community assembly mechanisms at increasing spatial scales within and between planktonic and benthic habitats. In addition, we leveraged the nested structure of our sampling design, evaluating patterns of diversity within the overall Lookout Creek watershed and within the nested sub-watershed, Watershed 01 (Fig. 1A).

Because species sorting was the dominant process detected across scales in the stream network, we examined the longitudinal variation in the magnitude and direction of species sorting. Specifically, we quantified how βNTI (used to infer selection) varied with habitat type (within sediments, within planktonic samples, and between habitats) and network position (headwater streams versus downstream) using ANOVA. The ANOVA model was constructed with βNTI values as the response variable, and with habitat type and network position as the factors. We included an interaction term to test whether the effect of network position on βNTI differs with habitat type. We then performed Tukey’s HSD test to evaluate significant differences among the factors in the model.

## RESULTS

### Patterns of α- and β-diversity

Planktonic and sediment-associated bacterial communities differed in α-diversity. On average, we observed 20% higher α-diversity in the bacterioplankton than in sediment-associated communities (species equivalents: 1789 ± 101 in sediments, 2210 ± 131 in plankton, *p* = 0.002, F_1,47_ = 10.28). Bacterioplankton also contained > 3-fold more habitat-specific taxa (i.e., taxa never found in sediment samples) than sediment-associated communities (20.5 ± 0.9% unique in planktonic taxa vs. 6.2 ± 0.7% unique sediment taxa, *p* < 0.001, F_1,47_ = 219.3).

Patterns of β-diversity suggest key differences in community structure within and between habitat types, across stream orders, and across spatial scales. Across the network, variation in bacterial community structure was explained primarily by the habitat from which the samples were taken (PERMANOVA, *R*^*2*^ = 0.12, *p* = 0.001), the stream order of the sampling site (*R*^*2*^ = 0.033, *p* = 0.002), and the spatial extent of the drainage basin (i.e., spanning the entire Lookout Creek watershed or the smaller, nested Watershed 01) where the samples were collected (*R*^*2*^ = 0.04, *p* = 0.004). Redundancy analysis (RDA) detected a separation between bacterioplankton and sediment samples along RDA1, which explained 12% of the variation (Fig. 2). Along RDA2, samples separated along a gradient that captured elevation and resource availability. Specifically, we identified communities that clustered in high elevation sites with relatively high dissolved organic carbon (DOC) concentrations and communities that clustered in low elevation sites with higher total phosphorus (TP), total nitrogen (TP), conductivity, and pH. Sites in Watershed 01 also clustered together along RDA2 more tightly than sites dispersed across the broader Lookout Creek watershed.

**Fig. 2.**
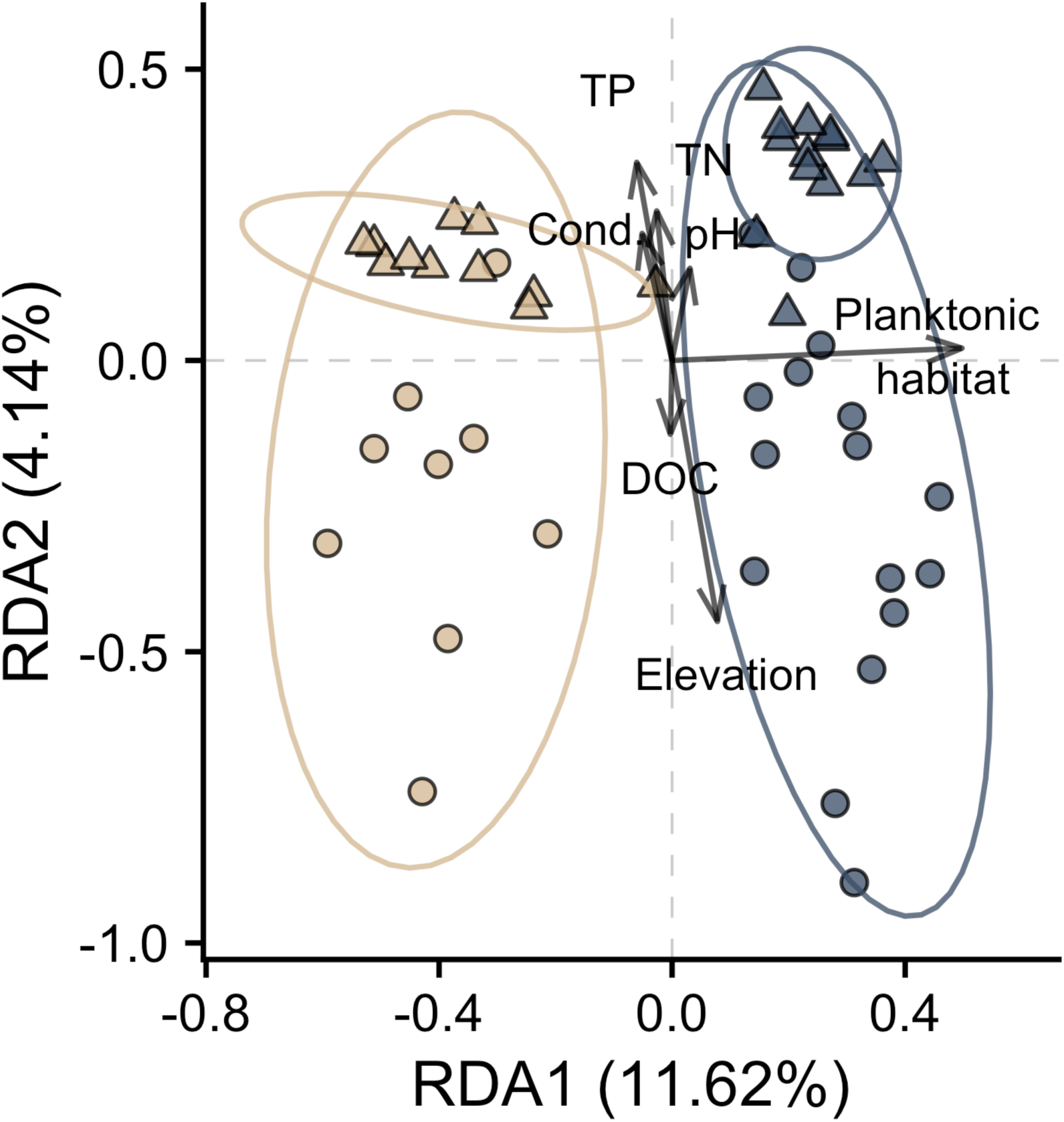
Taxonomic β-diversity revealed compositional differences among habitat types, but also within habitats. Redundancy analysis (RDA) found the primary axis of variation in community composition could be explained by habitat type (i.e., planktonic or sediment-associated). Within habitats, a secondary axis of variation explained a gradient from high elevation, low conductivity sites in the headwaters, to low elevation sites with high conductivity in the higher order streams. RDA2 also captured differences in spatial scale of sampling, with sites from Watershed 01 clustering together (triangles), nested within sites distributed across the broader Lookout Creek catchment (circles). Beige symbols indicate sediment-associated samples and blue samples indicate planktonic samples; circles indicate samples taken from the broader Lookout Creek catchment, while triangles indicate samples taken from the smaller Watershed 01 nested within Lookout Creek. Ellipses represent 95% confidence intervals for the group locations in the RDA subspace.

As expected, spatially isolated sites in the dendritic network were more compositionally dissimilar than nearby sites (Fig. 3). However, the rate of increase in dissimilarity was habitat-dependent and was more gradual in sediment than in planktonic communities. In contrast, community dissimilarity between different habitat types was consistently higher than within-habitat differences from local (y-intercept) to regional scales (∼ 10 km).

**Fig. 3.**
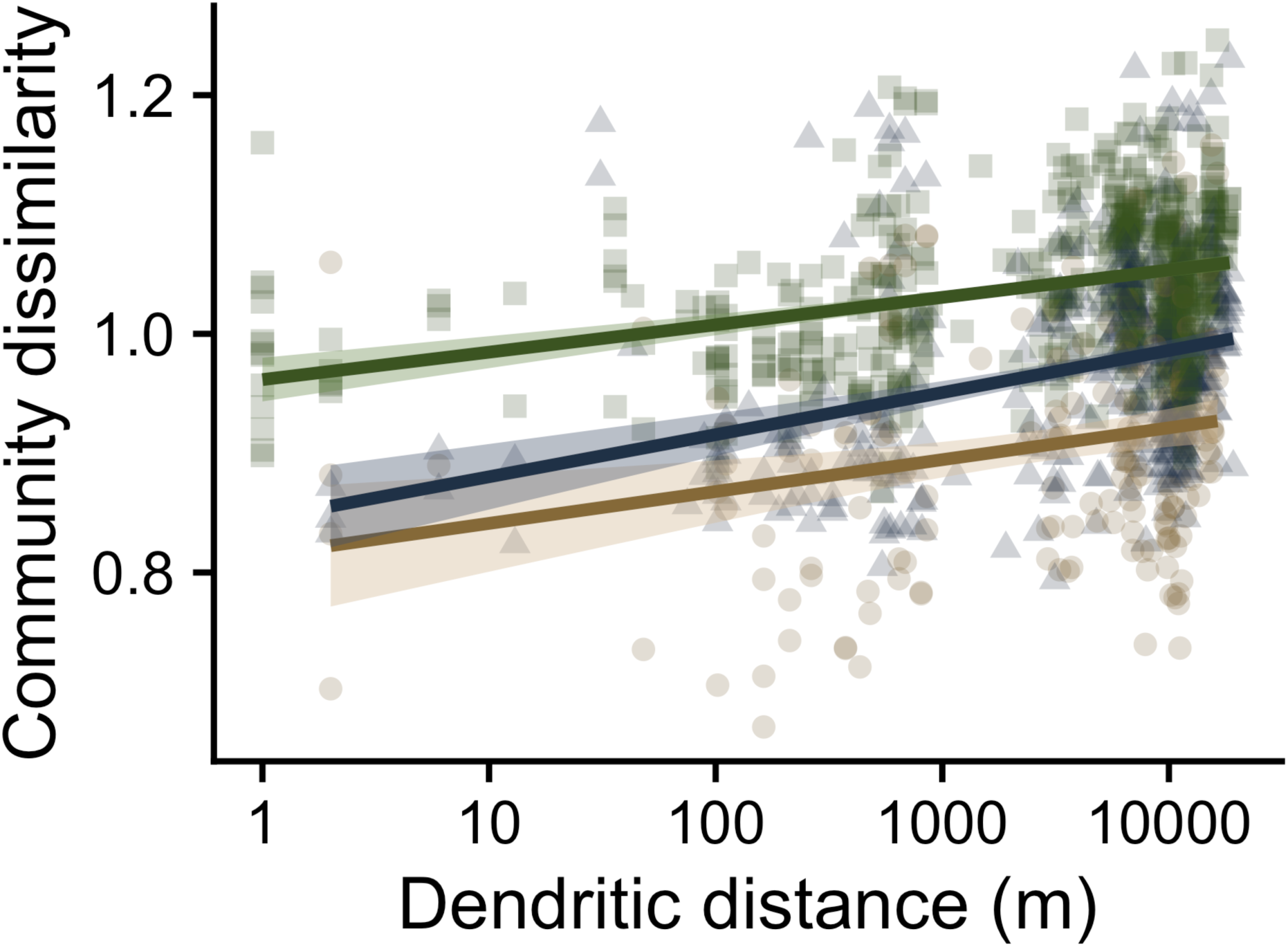
Community dissimilarity across spatial scales depended on habitat type. Community dissimilarity (using Hellinger distance) increases with dendritic distance in the network. Comparisons between sites in different habitat types (green squares, highest line) had the lowest community similarity at all spatial scales in the drainage basin. Comparisons within planktonic samples (blue triangles, middle line) were more similar at local spatial scales than between habitat comparisons (∼ < 1 km), but at broader spatial scales, their dissimilarity approached that of local-scale between-habitat dissimilarity. Comparisons within benthic samples (beige circles, lowest line) were most similar across all spatial scales and their dissimilarity increased the slowest with dendritic distance.

### Scale-dependent community assembly

Bacterial community assembly in the Lookout Creek stream network was habitat and scale dependent. Overall, hydrologically connected communities predominantly showed evidence of convergent or divergent species sorting (620/696 = 89% of comparisons), with some evidence for stochastic assembly (54/696 = 7.8% of comparisons) or dispersal limitation (18/696 = 2.6%) (Fig. 4). Detection of mass effects, except at small spatial scales, was comparatively low (4/696 = 0.6% of comparisons). Within communities of the same habitat type, convergent species sorting was the dominant process (sediments: 88/134 = 66%; plankton: 142/214 = 66%). Sediment communities showed strong signatures of convergent species sorting across all spatial scales in the catchment (1 m to 10 km), with divergent species sorting playing a relatively smaller role. Within sediments, mass effects were detected at local scales (< 10 m) and stochastic effects emerged at broader (> 1 km) scales. Planktonic communities also showed evidence for convergent species sorting, but we detected divergent species sorting (32/46 = 70% of comparisons > 1 km apart) and dispersal limitation (6/46 = 13% of comparisons) at broader spatial scales. Between communities in different habitats, divergent species sorting was the dominant assembly mechanism (294/348 = 84% of comparisons), with strong stochastic effects detected at smaller spatial scales (< 100 m).

**Fig. 4.**
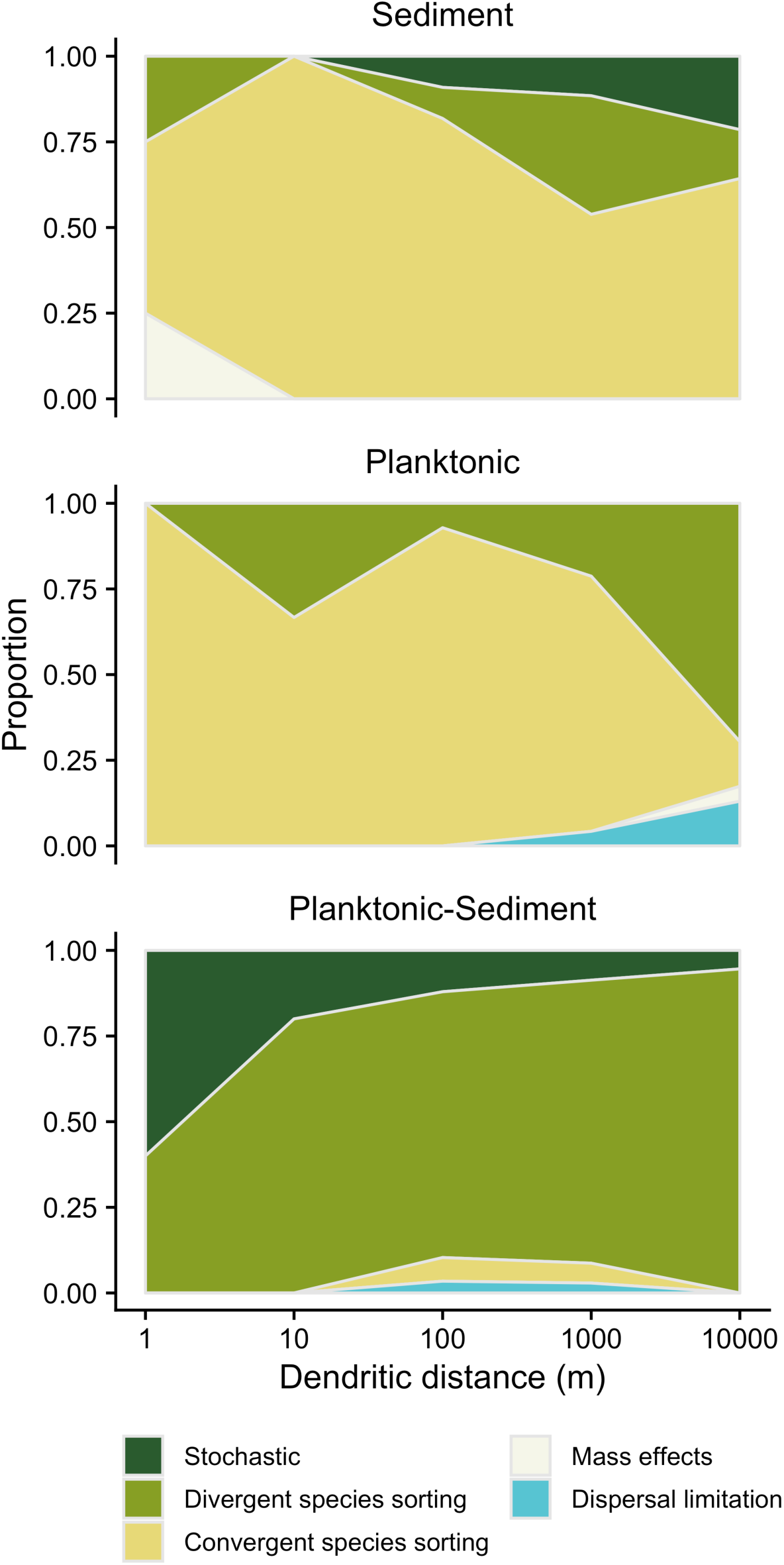
Habitat and scale dependent community assembly mechanisms in the dendritic metacommunity. In sediment habitats, convergent species sorting is the dominant assembly mechanism across all spatial scales. Divergent sorting and mass effects occur rarely at local scales, while divergent sorting and stochasticity are more common at larger scales. Within planktonic habitats, there is a transition from convergent species sorting to divergent species sorting with increasing spatial scale. Between planktonic and benthic habitats, divergent species sorting was the dominant mechanism inferred across most spatial scales, but stochasticity was common at local scales.

### Longitudinal trends in community assembly

Different positions in the dendritic network experienced variable strengths and directions of species sorting depending on habitat type. Because species sorting was the dominant process overall, we further investigated the direction (e.g., convergent or divergent) and magnitude (i.e., absolute value) of sorting inferred by βNTI along the longitudinal dimension of the stream network. We found habitat-specific trends in species sorting between headwater and downstream communities (Fig. 5). In the ANOVA model, network position, habitat, and the network position × habitat interaction were all significant terms explaining βNTI in the metacommunity (Table 1). Using Tukey’s test, we found that sorting in headwater communities was significantly divergent (i.e., |βNTI| > 2) for all comparisons within and between habitat types (mean ± SE βNTI: sediment: 8.32 ± 2.00, planktonic: 12.3 ± 1.32, planktonic-sediment: 15.2 ± 0.581). In contrast, sorting in downstream communities was significantly convergent among sediment communities (mean βNTI: −3.32 ± 0.054 SE), highly variable but stochastic on average among planktonic communities (βNTI: −1.12 ± 0.028 SE), and significantly divergent between communities in different habitats (βNTI: 11.2 ± 0.023 SE). Thus, our results suggest that the degree to which communities converge downstream depends on habitat type.

**Table 1.** ANOVA output for βNTI ∼ habitat * network position.

|  | Df | Sum Sq. | Mean S. | F value | p-value |
| --- | --- | --- | --- | --- | --- |
| *Network position* | 1 | 2562 | 2652 | 36.735 | $2.07 \times 10^{-9}$ |
| *Habitat* | 2 | 32831 | 16416 | 235.34 | $< 2 \times 10^{-16}$ |
| *Position* × *habitat* | 2 | 669 | 334 | 4.795 | $8.51 \times 10^{-3}$ |
| *Residuals* | 810 | 56499 | 70 |  |  |

**Fig. 5.**
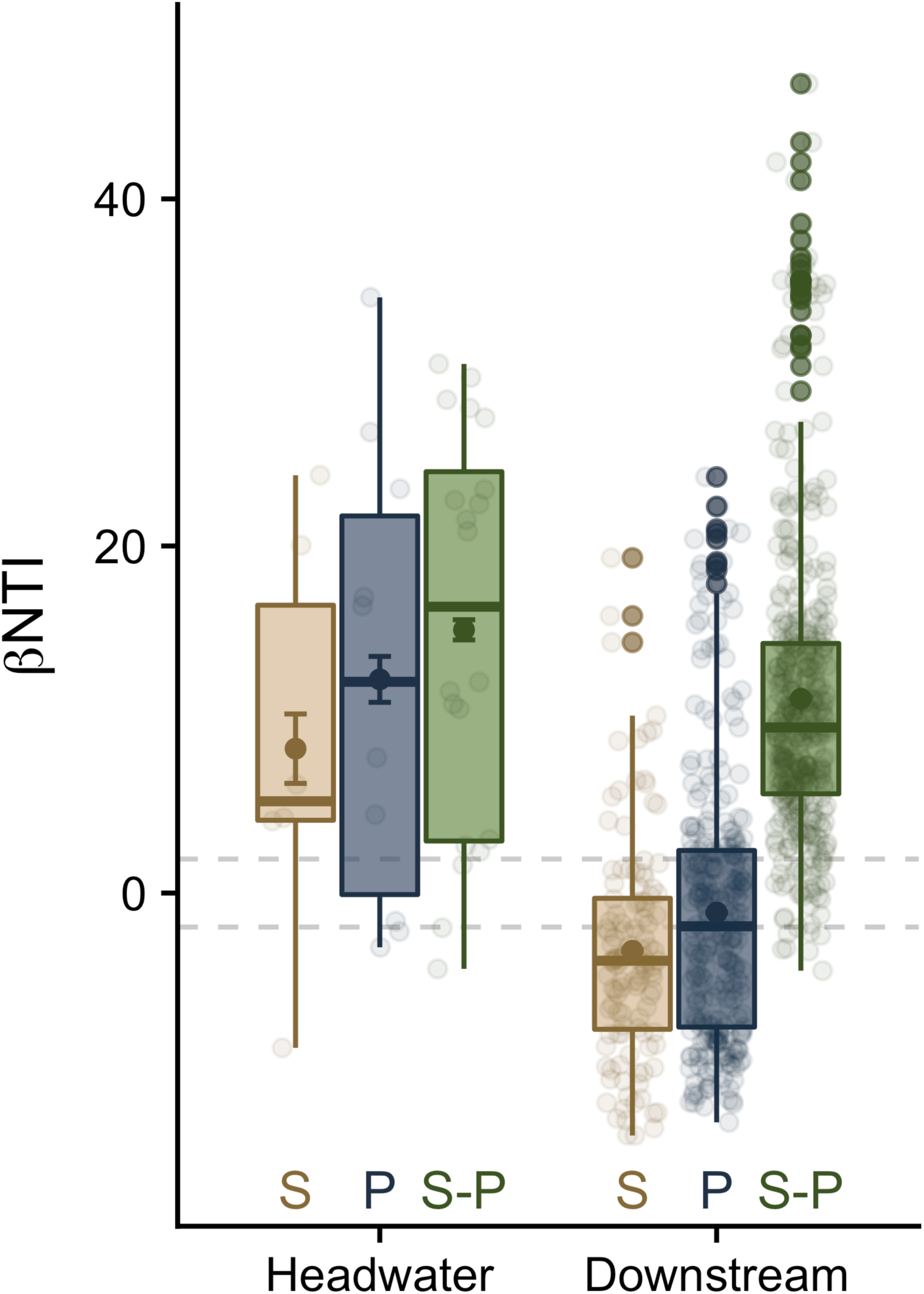
Deviations in phylogenetic β-diversity relative to null expectations (bounded by gray dashed lines) demonstrated longitudinal patterns in phylogenetic convergence or divergence between headwater and downstream sites. We compared phylogenetic β deviations within habitats (beige = sediment (S), blue = plankton (P)) and between habitats (green = sediment-plankton comparisons (S-P)). Points and error bars indicate mean ± SEM. Communities in headwaters were more phylogenetically dissimilar than expected by chance (i.e., βNTI values > 2) for all habitat comparisons, with sediment communities least divergent and between-habitat comparisons most divergent. Downstream communities showed contrasting patterns on average, with variation around the mean. On average, sediment communities were more phylogenetically convergent than expected by chance, while planktonic communities showed less deviation from stochastic expectations (mean βNTI < |2|). Downstream βNTI between habitats was lower than in headwaters, but still significantly positive on average.

## DISCUSSION

We have shown that bacterial community assembly in a dendritic metacommunity depends on vertical habitat structure, spatial scale, and network position. Overall, species sorting was the predominant assembly mechanism across the stream network, but the direction (e.g., convergent versus divergent) and magnitude (i.e., βNTI absolute value) of species sorting varied across space. Divergent species sorting maintained compositionally distinct planktonic and sediment communities (Fig. 2-5), but stochastic assembly occurred at local scales (< 1 km). Within planktonic or benthic communities, convergent species sorting was the dominant assembly process (Fig. 4). However, sediment-associated communities also showed evidence of local-scale (< 10 m) mass effects and broad scale (> 1 km) stochasticity, while planktonic communities transitioned from convergent to divergent species sorting as scale increased (Fig. 4). In the longitudinal dimension of the network, we detected the strongest signal of divergent sorting among headwater communities of both habitat types, while convergent sorting was most evident in downstream sediment-associated communities (Fig. 5). Thus, community assembly in dendritic metacommunities is strongly habitat- and scale-dependent, which may help reconcile taxonomic differences in dendritic metacommunity organization through tighter integration of spatial scale and vertical habitat structure.

### Compositionally distinct planktonic and sediment-associated communities

We found several lines of evidence that planktonic and sediment-associated communities are compositionally distinct due to deterministic processes. First, the higher α-diversity and greater proportion of habitat-specific taxa detected in the plankton suggest that many planktonic taxa do not successfully colonize the streambed. Conversely, these patterns also suggest that planktonic diversity is not maintained by seeding from highly diverse sediment communities, perhaps because sources other than benthic sediments also contribute to planktonic diversity (Battin et al. 2016). Across the watershed, community structure was consistently different between planktonic and sediment habitats (Fig. 2), similar to what has been reported for stream bacterial communities in alpine (Besemer et al. 2012, Wilhelm et al. 2013) and arid (Kaestli et al. 2019) ecosystems. Furthermore, spatial patterns of community dissimilarity show that local-scale differences between planktonic and benthic communities can exceed within-habitat differences at larger spatial scales (Fig. 3). Such differences may be due to the increased stability of the sediment habitat matrix relative to the water column, as well as the physiochemical environmental differences between the two habitats (Hermans et al. in press). In light of these results, our inferred community assembly processes (Fig. 4) support a prevailing role for divergent species sorting between planktonic and sediment-associated communities.

The strength of divergent species sorting between communities in different habitats was scale dependent (Fig. 4). At local scales, differences in community structure were partly due to stochastic processes, while divergent sorting played an increasingly large role at broader spatial scales. This scale dependence may arise from variable dispersal kernels (e.g., along preferential flow paths) or from vertical hydrological exchange at the stream reach scale, which could generate idiosyncratic spatial variation in community structure. In our study system, it has been shown that vertical hydrological exchange plays a more important role in headwaters than in downstream reaches (Ward et al. 2019), suggesting that vertical fluxes may be responsible for disrupting the species sorting process predominantly in the headwater reaches. If so, stochastic dispersal may increase in importance downstream as channels widen and the relative importance of vertical exchange diminishes. At broader spatial scales, divergent species sorting between planktonic and benthic communities is strong enough to overcome local-scale stochasticity. Thus, although most studies have examined diversity patterns within either planktonic or sediment communities separately at large scales, or more intensively at small spatial scales, here we show that species sorting influences community structure in both habitats in a scale-dependent way.

### Longitudinal and scale-dependent transitions in planktonic community assembly

We found mixed support for the expectation that bacterioplankton community assembly is driven primarily by dispersal. On the one hand, the positive relationship between dendritic distance and community dissimilarity could result from dispersal, such as local-scale mass effects or regional-scale dispersal limitation, but it could also reflect species sorting along divergent environmental conditions in the watershed (Soininen et al. 2007). Divergent species sorting would be consistent with the phylogenetic patterns we observed at large spatial scales (Fig. 4). As previously suggested, immigration from terrestrial ecosystems can also contribute to bacterial diversity in streams, particularly in headwater reaches (Read et al. 2015, Savio et al. 2015, Ruiz-González et al. 2015). However, in some ecosystems, dispersal connectivity between terrestrial soils and bacterioplankton may be weak and transient (Hermans et al. in press). In our study, planktonic communities transitioned from high phylogenetic β-diversity among headwaters to lower phylogenetic β-diversity downstream, and from convergent species sorting at the reach scale (< 1 km) to divergent sorting at the watershed scale (1-10 km). Longitudinal diversity patterns reflect environmental gradients from high to low elevation sites (Fig. 2) that may relate to the environmental filters that influence species sorting on drifting bacterioplankton (Figs. 4-5). Thus, the water column may serve as a dispersal corridor for terrestrial-derived bacteria that progressively undergo species sorting as they drift downstream.

Contrary to our expectations, we did not detect a strong signal of mass effects in the bacterioplankton communities (Fig. 4). We attribute this to the fact that mass effects may be difficult to distinguish from convergent species sorting without direct knowledge of dispersal rates because inferences of mass effects based on taxonomic homogenization (*β*_*RC,Hellinger*_ < −.95) would also homogenize phylogenetic diversity (βNTI < –2). We did, however, detect dispersal limitation at the largest spatial scales (e.g., from 1-10 km), likely due to the large spatial distances between high- and low-elevation sites. For example, low-elevation headwaters of Watershed 01 were tightly clustered within the range of communities spanning the broader Lookout Creek (Fig. 2), which may reflect that fact that some high-elevation taxa are dispersal-limited with respect to colonizing Watershed 01 and vice versa. Thus, despite low power to detect mass effects, our results suggest that terrestrial-derived bacteria, environmental gradients, and dispersal limitation may explain changes in planktonic diversity across spatial scales and from headwaters to downstream reaches of the network.

### Sediment community assembly shows longitudinal trends in direction despite weaker scale-dependence

In the sediment-associated communities, our results suggest species sorting may be a dominant process across a range of spatial scales (Fig. 4). First, sediment-associated communities were distinct from planktonic communities across the catchment (Fig. 2), consistent with convergent species sorting favoring the colonization of a subset of taxa from the overlying water column. Sediment communities also showed weaker scale-dependence in community structure, as they remained more similar to each other with increasing dendritic distance than planktonic communities did (Fig. 3), potentially due to similar environmental filters acting across the stream network. Indeed, convergent species sorting was identified as the dominant assembly mechanism across all spatial scales of comparison (Fig. 4), suggesting that, in general, sediment communities consist of phylogenetically similar taxa favored by environmentally similar environments.

However, species sorting was not always convergent in sediments across the metacommunity. In particular, we observed greater phylogenetic β-diversity than expected under purely stochastic assembly among headwater sites. This significant divergent species sorting suggests that, despite local-scale convergence within reaches (i.e., similar communities assemble within reaches, regardless of network position), different headwaters favor the assembly of phylogenetically distinct sediment communities (Fig. 5). This divergence among headwaters may reflect dissimilar resource inputs among headwaters draining different terrestrial areas, or spatial variation in terrestrial sources that contribute to stream sediment assembly. The transition to convergent species sorting downstream may represent longitudinal gradients in microhabitat structure (e.g., sediment size) and resource complexity (e.g., allochthonous vs. autochthonous organic matter) from lower-to higher-order streams (Vannote et al. 1980). Interestingly, we found evidence for local mass effects (e.g., 1-10 m) in the sediment communities (Fig. 4), which may be due to high hydrologic conductivity that mobilizes fine sediments their attached bacterial communities. The increasing frequency of stochastic assembly processes observed at the largest spatial scales (1-10 km) could reflect the idiosyncratic effects of disturbance history (e.g., large floods, debris slides, logging) that are common across the Lookout Creek watershed (Swanson and Jones 2002). Thus, while divergence is common in both sediment and planktonic communities among headwaters, sediment communities show weak scale-dependence, likely due to a more consistent set of environmental filters across spatial scales.

### Multi-layer dendritic metacommunities

Our work provides an empirical demonstration that the community assembly processes structuring metacommunities in dendritic networks vary not only with network position, but also across spatial scales and along the vertical dimension of streams, which encompasses planktonic and benthic habitats. The joint consideration of spatial scales and vertical habitat structure may be crucial to resolving taxonomic differences in diversity patterns in dendritic metacommunities (Schmera et al. 2018). For example, aquatic taxonomic groups (e.g., riparian plants, benthic invertebrates, and microorganisms) in dendritic networks span a wide range of body sizes and generation times, disperse via different dispersal corridors throughout the stream network, and occupy benthic and planktonic habitats in vastly different ways. These key differences suggest the potential for a broader synthesis of metacommunity dynamics in stream networks built on a revised perspective embracing multi-layer dendritic networks with varying rates of dispersal and habitat use in the vertical and longitudinal dimensions.

## ACKNOWLEDGEMENTS

We thank Adam Ward for logistical and financial support during sample collection. This work was supported by the National Science Foundation (DEB-1442246 to JTL), US Army Research Office Grant (W911NF-14-1-0411, J.T.L.), and the Department of Biology at Indiana University (George W. Brackenridge Fellowship, Louise Constable Hoover Fellowship to NIW). Data and code for the project can be found at NCBI (BioProject XXXXXXX) and GitHub (https://github.com/LennonLab/HJA-streams). LIDAR Data and facilities were provided by the HJ Andrews Experimental Forest and Long-Term Ecological Research program, administered cooperatively by the USDA Forest Service Pacific Northwest Research Station, Oregon State University, and the Willamette National Forest. This material is based upon work supported by the National Science Foundation under Grant No. DEB-1440409.

